# Physical laws predict methane hotspots in global mountain waters

**DOI:** 10.64898/2026.07.31.740466

**Authors:** Dan Zhu, Irfan Rashid, Katey Walter Anthony, Shujuan Tong, Nabin Bhattarai, Xin Zou, Srijana Joshi, McKenzie Kuhn, Jianliang Liu, Haibo Jiang, Huai Chen, Ning Wu

**Affiliations:** Chengdu Institute of Biology, Chinese Academy of Sciences, Chengdu 610041, China; Zoige Wetland Ecosystem Research Station, Chinese Academy of Sciences, Hongyuan 624400, China; Department of Botany, University of Kashmir, Srinagar, J&K, 190006, India; Water and Environmental Research Center, University of Alaska Fairbanks, Fairbanks 755860, Alaska, USA; International Centre for Integrated Mountain Development, Kathmandu GPO Box 3226, Nepal; Department of Geography, University of British Columbia, Vancouver V6T 1Z4, BC, Canada

**Author notes:** Corresponding author: Dan Zhu, Haibo Jiang, Huai Chen, Ning Wu, Corresponding author’s emails, and; 00862882890288.

**Keywords:** mountain aquatic ecosystem, atmospheric pressure, methane-containing bubbles, positive feedback, climate change mitigation

## Abstract

Accurate accounting of aquatic methane emissions is critical for climate change mitigation, yet current global budgets overlook a key driver: the elevation-regulated atmospheric pressure. Here, we present the first large-scale investigation of methane ebullition across 164 shallow waters spanning elevations from sea level to 4886 meters. We demonstrate that ebullition rate increases with elevation-over four times higher at >3000 m a.s.l. than at sea level-due to two synergistic, pressure-dependent physical mechanisms: a ‘degas’ effect (enhanced bubble formation) and a ‘trigger’ effect (facilitated bubble ascent). Independent theoretical prediction of the combined effects shows near-perfect agreement with the empirical elevation trend, quantitatively confirming that these physical mechanisms are the primary drivers of enhanced ebullition at high elevations. Our findings reveal that mountain aquatic ecosystems represent unaccounted methane hotspots that have been systematically underestimated in global inventories.We therefore call for urgent integration of these ecosystems into IPCC assessments and targeted mountain mitigation and sustainable management strategies.

## Introduction

Methane, responsible for approximately 23 % of current global warming^1^, is a focal point of international climate agreements such as the Global Methane Pledge. Methane emissions from aquatic ecosystems constitute approximately half of the global methane budget^2^. However, global estimates of methane emissions from these ecosystems are fraught with uncertainties, largely due to the imprecise quantification of the relative contributions of various methane transport pathways^3,4^. Among these pathways, ebullition-characterized by the release of methane-containing bubbles from sediments to the atmosphere-is recognized as the most efficient yet sensitive one for methane emission from anoxic sediments^5,6^.

A critical determinant of the ebullition process is the pressure exerted on the sediments, whether from atmospheric or hydrostatic sources^7^. This pressure influences bubble formation and their ability to overcome the sediment resistance, ultimately affecting their release into the atmosphere^8–11^. Over the past decades, studies have identified decrease in local atmospheric pressure, such as those preceding cold fronts, or water level drawdowns, as triggers for sporadic but significantly enhanced methane ebullition (up to approx. 840 mg CH_4_ m^-2^ d^-1^) events from diverse aquatic bodies^12–19^. Shallow waters-lakes, ponds, and wetlands with water depth less than four meters-contribute at least 60% of global aquatic methane emissions^2,20^, and are particularly sensitive to pressure changes because their limited water column minimizes hydrostatic buffering^3,12^. Given that atmospheric pressure declines systematically with altitude^21^, shallow waters in mountain regions may experience persistently enhanced ebullition, yet this spatial dimension remains largely unexplored.

Evidence from high-elevation lakes and streams on the Tibetan Plateau suggests a clear association between reduced atmospheric pressure and elevated ebullition^22,23,24^. Similar patterns have been observed in mountain streams elsewhere^25,26^. However, other factors complicate this picture. Substrate availability typically decline with elevation^27^, and high salinity can further suppress methane production in some mountain waters^28^. Whether, and to what extent, these opposing influences weaken the stimulatory effect of elevation has remained unclear at the global scale.

The question then arises: can the systematic decline in atmospheric pressure with elevation serve as a predictive basis for methane ebullition across the world’s shallow waters? If so, how do the underlying mechanisms regulated the ebullition along elevation? By integrating a large-scale elevational transect with physical laws, we address both these questions and provide a quantitative framework that bridges field observation with theoretical modelling, advancing our understanding of the spatial patterns and controls of global methane emissions.

## Results

### Methane ebullition along the elevation gradients

We synthesized data from 164 shallow water ecosystems, with elevation ranging from sea level to 4886 meters above sea level (m a.s.l.) (Figure 1a)^22,24,29–31^. To ensure data comparability, the lakes were selected from regions with relatively uniform climates, and the reported methane flux rates represent the mean values from extended time-series monitoring conducted during the ice-free season (Table S1). The dataset was incorporated with our field observations from a mid-elevation lake in the central Himalayas (Extended Data Fig. 1). Given the variability in mean water depth across these lakes (Table S1), we calculated both hydrostatic and local atmospheric pressures for each site (see Methods). Typically, the contribution of hydrostatic pressure to the total pressure diminishes with increasing elevation in our study, with most of the observed changes in total pressure being due to elevation-related variations in atmospheric pressure (Fig. 1b).

**Figure 1.**
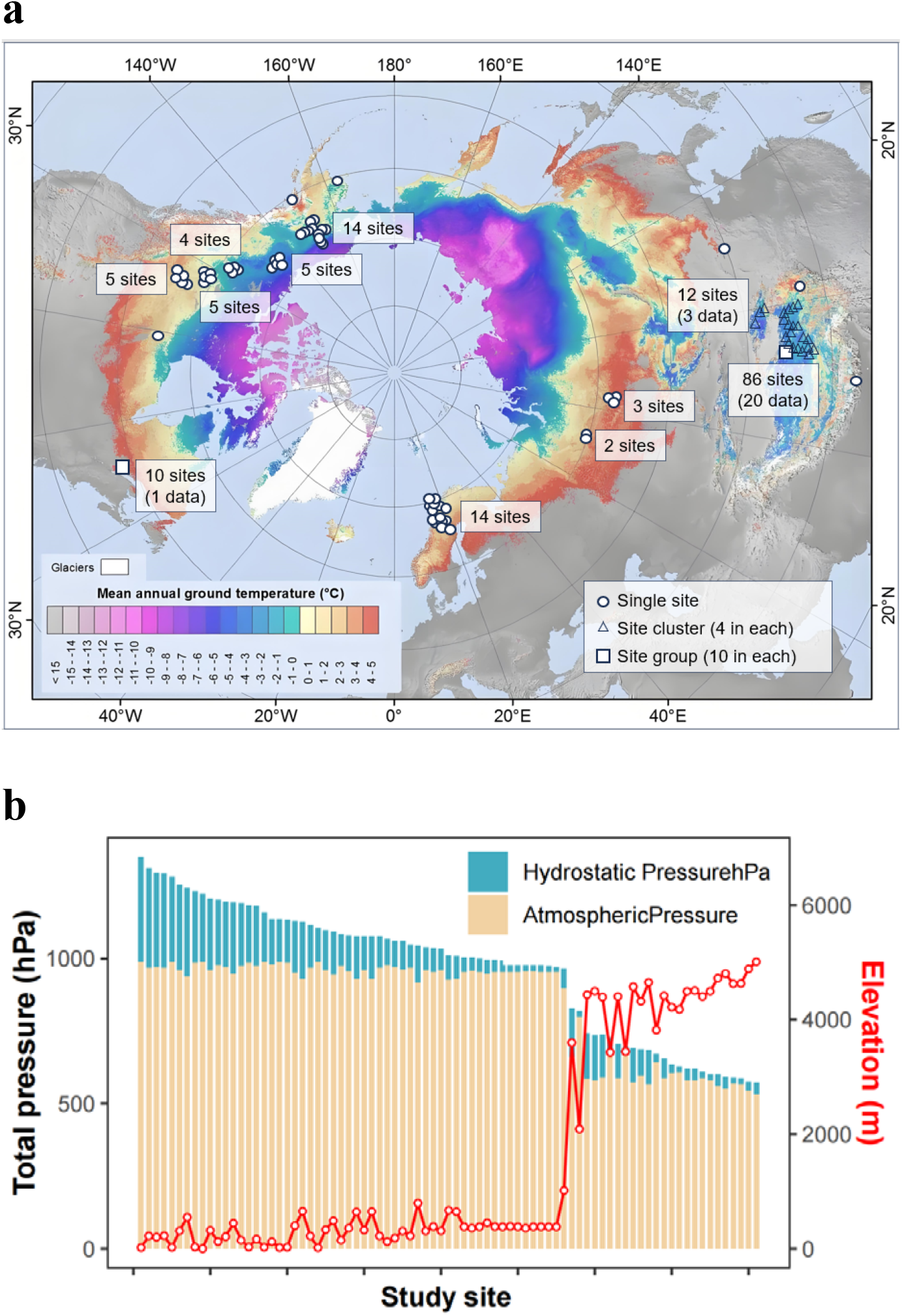
The shallow waters for characterizing the relationship between methane ebullition and atmospheric pressure. **(a)** The geographical distribution of the 164 shallow waters (overlapped sites are due to data availability in the respective study) (the map is adapted from Obu et al. (2019) (ref. 56)). **(b)** The total pressure on the sediments of each site, which is composed by their local atmospheric pressure (calculated from the elevation, see Methods) and hydrostatic pressure (calculated from the mean depth of water column).

Our analysis demonstrated that methane ebullition rates increase significantly under low pressure conditions compared to higher pressure conditions (Fig. 2a). In contrast, diffusive rates did not show significant variation across different pressure groups (Extended Data Fig. 2). The relationship between methane ebullition rates and pressures across all lakes was well-characterized by a linear model (*R*^2^=0.38, *P*<0.001), indicating a pronounced increase in methane ebullition with rising elevation (Figure 2b). However, methane ebullition showed no significant correlation to water temperature (*P* = 0.677).

**Figure 2.**
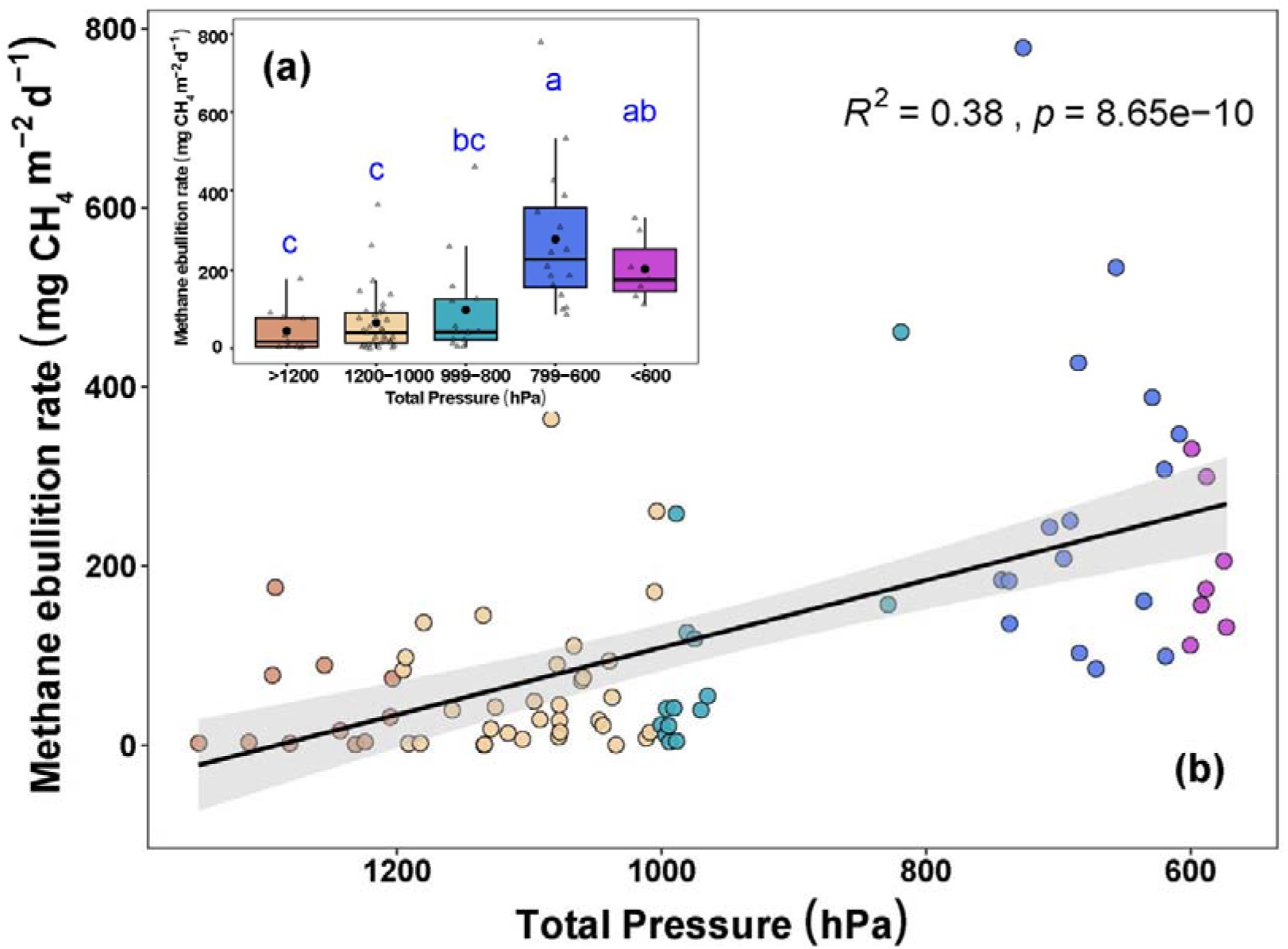
The relationships between methane ebullition and pressure. **(a)** Comparison of the ebullition rates by grouped total pressures Data presented are first quartiles, medians and third quartiles with standard errors. **(b)** The linear fitting of point data between methane ebullition rate and total pressure. Different colors of points represent five total pressure levels. Regression coefficients and P values are given.

### Comparison of ebullition parameters between high and low elevations

To further elucidate the behavior of methane-containing bubbles, we collected samples from one lake and four ponds in high-elevation (see Methods) and compared them with samples from lakes of various elevations. The pressure composition of this gradient is the integration of atmospheric and hydrostatic pressure, which is similar to the previous one for the regression analysis. Some apparent trends of the parameters along the gradient could be observed. Along the rising elevation, the methane concentration in bubbles decreased dramatically, while the occurrence of ebullition, the rate of methane ebullition and the contribution was found to increase. Our comparative analysis indicates that lakes with lower pressure, despite having lower methane concentrations in their bubbles, exhibited more rapid gas ebullition, leading to higher and more predominant methane ebullition (Table 1, Extended Data Table 1).

**Table 1.** Characteristic of summertime methane ebullition from shallow waters along the elevational gradients.

| Site | Country | No. of lakes/<br>Ponds | Elevation,<br>m a.s.l. | Water<br>depth, m | Method | Occurrence,<br>% | Methane<br>concentration in<br>the bubble<br>mean (range), % | Contribution to<br>the total<br>flux, % | Reference |
| --- | --- | --- | --- | --- | --- | --- | --- | --- | --- |
| Lakes in NE<br>Siberia | Russia | 2 | <200 | <4.0 | BT | nd | (55.0-63.8) | 72.4<sup>a</sup> | 41, 57 |
| Lakes near<br>Abisko | Sweden | 3 | 380 | <4.0 | BT | 45 | 34.8 (0.0-98.6) | 40<sup>b</sup> | 57, 58 |
| Lakes<br>in Wisconsin | US | 11 | 509-557 | <4.0 | BT/IS | 25-80 | nd | 40-60 | 12 |
| Ponds in<br>Quebec | Canada | 10 | 244-671 | <2.0 | BT, CH | nd | 57.6 (1.3-97.0) | 56 | 13 |
| Lake<br>Windsborn | Germany | 1 | 497 | <1.5 | BT | nd | (0.0-86.4) | nd | 59 |
| Lake<br>Ulansuhai | China | 1 | 1018 | 0.7 | BT/CH | 52 | nd | 69 | 31 |
| Lake Dhaap | Nepal | 1 | 2089 | 0.2 | CH | 25 | nd | 88<sup>c</sup> | This study |
| Thermokarst<br>lakes | China | 88 | 3500-4500 | <2.0 | CH | 100 | nd | 84 | 24 |
| Lakes on the<br>Zoige Plateau | China | 5 | 3430 | <0.5 | BT/CH | 74 | 9.4 (5.9-15.4) | 93 | 22, This study |

Method: BT-Bubble trap, IS-Ice survey, CH-Chamber.
<sup>a, b</sup>: calculated with total flux from thermokarst lakes and peatland lakes in Ref. 57, respectively, <sup>c</sup>: plant mediated flux was excluded as described in Table S1.
nd: no data.

### Modelling bubble formation and movement

Our theoretical modelling demonstrates a nearly linear relationship between gas solubility and pressure (Fig. 3a and Equation 1, Methods). For example, in a shallow lake (water depth of 0.5 m) at an elevation of 2000 m a.s.l., where the total pressure (atmospheric pressure plus hydrostatic pressure) is 82% of that at sea level, there is a 18% increase in the availability of methane in the free gas phase. At 4000 m a.s.l., this increase is projected to be 32%, with solubilities of 1.19, 1.44, and 1.76 corresponding to 4000, 2000, and 0 m a.s.l., respectively, at a temperature of 12°C (referred to as points A, B, and C, respectively). This suggests a significant disparity in the methane withholding capacity of pore water along elevations.

**Figure 3.**
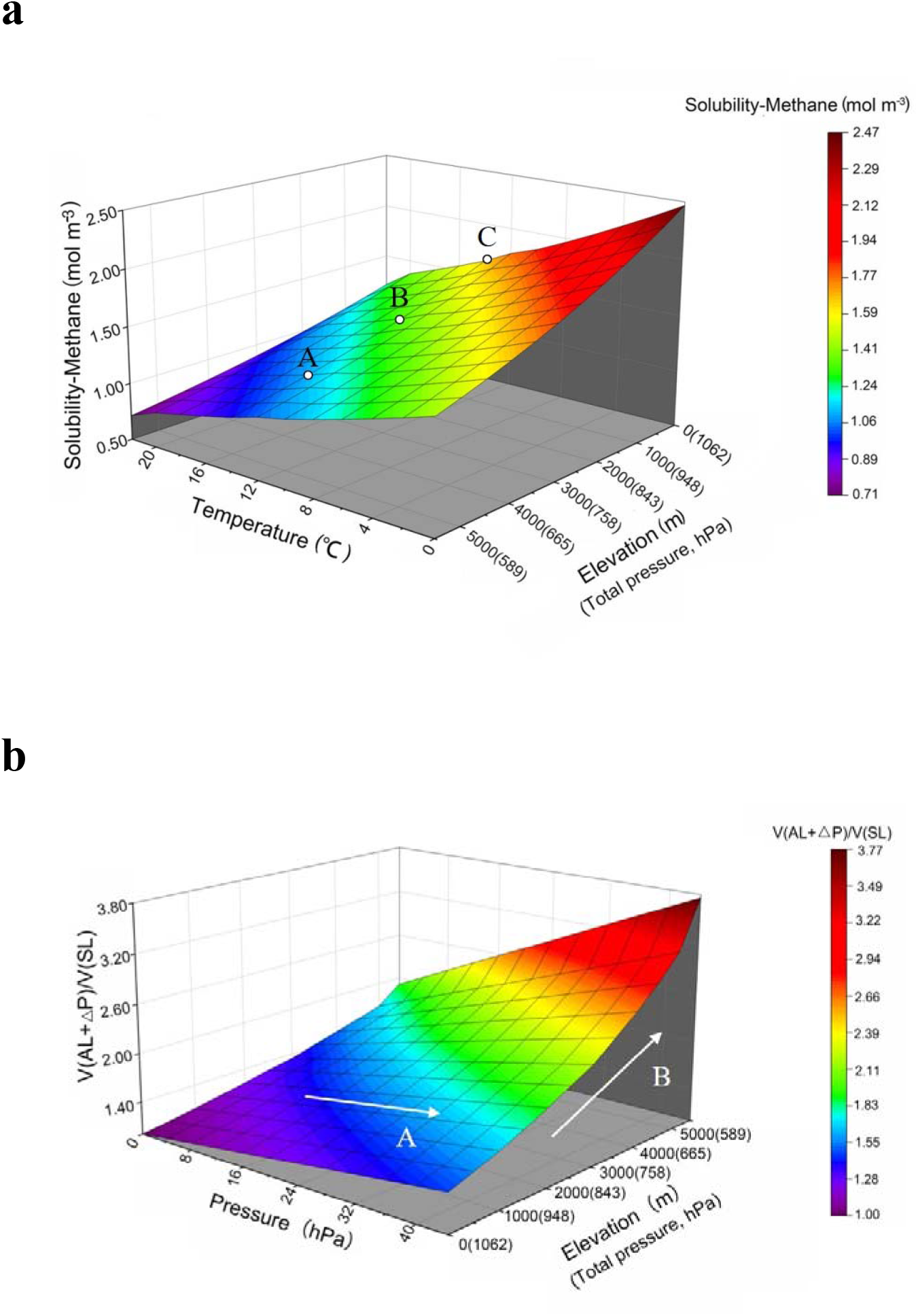
Modeling on the bubble formation and expansion in sediment/water (under 0.5 m water column) according to the Free Gas Law and Henry’s Law. (**a**) The modeled relationship between methane solubility versus atmospheric pressure and temperature. (**b**) The modeled relationship between gas volume and atmospheric pressure varies in both temporal and elevational scale (see Extended Data).

According to Henry’s law, the volume of free gas, for a constant mass, will expand under reduced pressure. To quantitatively assess the impact of pressure changes on bubble volume, we conducted additional theoretical modeling that accounts for the spatio-temporal variations (see Methods). Our model indicates that, similar to the natural temporal changes, the volume of a given mass of gas will increase with declining atmospheric pressure along rising elevation (i.e. directions indicated by Arrow A versus Arrow B in Fig. 3b).

### Theoretical versus empirical ebullition

The point-by-point comparison revealed a remarkably strong correlation between the two datasets (Table 2, *R*^2^=0.998, *P*<0.001). The fitted relative fluxes were slightly higher than the theoretical calculations (see Methods) at all elevations above 500 m a.s.l., with deviations ranging from +14% at 500 m a.s.l. to a maximum of +30% at 2000 m a.s.l., and then gradually converging to near parity at 6000 m a.s.l. (+0.3%).

**Table 2.** Comparison between the theoretical model and the empirical trend.

| Elevation<br>(m a.s.l.) | Total pressure<br>(hPa) | Fitted flux<br>(linear model)<br>(mg m<sup>-2</sup> d<sup>-1</sup>) | Fitted relative<br>flux (linear<br>model) | Theoretical<br>relative<br>flux | Relative<br>deviation (%) |
| --- | --- | --- | --- | --- | --- |
| 0 | 1062 | 86.0 | 1.00 | 1.00 | +0.0 |
| 500 | 1003 | 108.8 | 1.27 | 1.11 | +14.0 |
| 1000 | 948 | 130.0 | 1.51 | 1.23 | +23.0 |
| 1500 | 894 | 150.0 | 1.74 | 1.36 | +28.0 |
| 2000 | 843 | 167.7 | 1.95 | 1.50 | +30.0 |
| 2500 | 795 | 184.0 | 2.14 | 1.66 | +29.0 |
| 3000 | 758 | 196.3 | 2.28 | 1.82 | +26.0 |
| 3500 | 706 | 214.4 | 2.49 | 2.03 | +23.0 |
| 4000 | 665 | 229.2 | 2.66 | 2.25 | +18.0 |
| 4500 | 626 | 243.3 | 2.83 | 2.49 | +14.0 |
| 5000 | 589 | 256.4 | 2.98 | 2.75 | +8.0 |
| 5500 | 554 | 268.5 | 3.12 | 3.00 | +4.0 |
| 6000 | 520 | 280.2 | 3.26 | 3.25 | +0.3 |

## Discussion

### High ebullition driven by low pressure

To our knowledge, this study presents the first large scale elevational transect, comprising 81 data points, to explore the relationship between methane ebullition and atmospheric pressure. The early records of extreme high methane ebullition induced by pressure drops were of temporal variation. Compared to its natural temporal variation, atmospheric pressure changes more dramatically with elevation.

Interestingly, despite the range of atmospheric pressure fluctuations recorded at lower elevations being an order of magnitude less than that of the elevation gradient, the resultant methane ebullition events^32,33^ were comparable to those observed along elevations in terms of the strength (To draw a parallel with temporal variations, we have synthesized the data into a scatter plot, see Figure S3). However, compared to the “triggered” ebullition during the weather events observed in the low elevations, it should be noted that the elevated methane ebullition on the high elevations are constant, which is reflected by the seasonal averaged ebullition data collected from these continuous monitoring (Table S1). The moderate *R*^2^ value (0.38) reflects the inherent variability of ebullition processes and the influence of site-specific factors (sediment properties, organic matter quality, microbial community structure) that modulate the pressure effect^12,13^.

In our study, the pressure gradient along elevation was determined jointly by local atmospheric pressure and the hydrostatic pressure of the water column as illustrated in Fig. 1. This gradient actually benefited from variations in water column depth among sites, although the contribution of water column depth to the overall variation was much less than that of atmospheric pressure. Consequently, the observed increase in methane ebullition with elevation is confounded by the potential effect of water column depth. Under the ideal scenario where total pressure is solely determined by the atmosphere (i.e., treating hydrostatic pressure as equivalent to the effect of elevation at each site), we found that mean ebullition rates at elevations above 3000 m a.s.l. (corresponding to approximately <740 hPa) were more than four times higher than those at sea level (254.3 and 59.3 mg CH_4_ m^-2^ d^-1^, respectively). Although atmospheric and hydrostatic pressures exert equivalent control over bubble formation and transport in sediments, we acknowledge that data coverage remains insufficient at mid to high elevations (e.g., 1000 to 3000 m a.s.l.) on a global scale. Furthermore, high-elevation sites in our dataset are concentrated in the Tibetan Plateau and Himalayas (Fig. 1a), with limited representation from other major mountain systems such as the Andes, Rocky Mountains, and East African Highlands. This geographic bias may influence the global generalizability of our elevation-ebullition relationship.

In addition, as the ebullition data in this study was compiled from various independent studies, there are methodological biases in our analysis. For example, compared to the direct sampling on methane ebullition by the bubble traps, chamber method can leads to either underestimation or overestimation. We attempted to minimize these biases by selecting studies with extended monitoring periods during ice-free seasons (Table S1), but methodological differences remain a source of uncertainty in our absolute flux estimates. Nevertheless, the consistent directional trend (increasing ebullition with elevation) across studies suggests a convincing pattern despite methodological heterogeneity.

### The role of methane production

Our study has delineated the spatial pattern of methane ebullition in shallow waters across a broad elevation gradient. Unlike the methane ebullition “triggered” by atmospheric pressure drops in temporal scale as it is attributed to the instantaneous release of stored methane in the pore water, it remains to be determined whether variations observed at the elevation gradient can be exclusively ascribed to atmospheric pressure changes. At a global scale, elevated methane ebullition is also observed in aquatic ecosystems with favorable temperatures, such as in tropical or subtropical regions^34,35^, or in those experiencing substantial organic matter input from the terrestrial environment at low elevations^36,37^. In these cases, however, an increase in diffusive flux rate is anticipated, as both enhanced ebullition and diffusive flux are driven by robust methanogenesis^5,38^.

Contrary to these expectations, our findings showed no increase in diffusive rates in tandem with ebullition rates along the gradient of decreasing pressure. The Global River Methane Database (GRiMeDB) that documents methane dynamics in global rivers and streams has illustrated this difference between high and low elevations as well^39^. In lowland, high methane ebullition tend to be found in sites which have high diffusive fluxes, and this can be attributed to a substantial (particular human induced) substrate supply. On the contrary, high diffusive fluxes were absent in high elevations where high ebullitions were observed (Extended Data Fig. 5 a,b). This inference is also supported by methane diffusive data from lakes of high-elevation regions^23^.

Indeed, the methane production potential at high elevations has been found to be lower than at lower elevations under similar climatic conditions, due to distinct methanogen community structures^40^. Furthermore, the potential for insufficient substrate supply at high elevations, as evidenced by the mineral type of the sediments in some lakes (Lake bottom in Table S1), may exert a negative impact on ebullition rate. Regarding the source of methane, extraordinarily high ebullition rates recorded from Alaskan Yedoma lakes have been attributed to the utilization of ancient organic carbon (over ten thousand years old)^41^. However, this does not apply to high-elevation sites of this study, which generally have lower carbon content in their sediments^24^ (Table S1).

At the global scale, the methane production in high elevations may act as inhibitor, instead of facilitator, for ebullition in certain circumstances, as low substrate and high salinity would be predominant factors perplexing the effect of low atmospheric pressure^27,28^. And this could explain why high ebullition rates can’t always be found in aquatic ecosystems and wetlands in high elevations. In other words, the observed pattern in our study represents a central tendency, not an absolute rule, as local factors can override the pressure effect. We therefore suggest direct measurements of methanogenic activity across the elevation transect which can help resolve the relative contributions of production versus physical release in future studies.

### The physical controls on ebullition

It has already been proven that, by invoking Henry’s law and Ideal Gas Law, variations in methane exchange between pore water and gas phase under changes in both temperature and pressure can be explained^42,43^. Declining methane concentration along elevation which has been found in global rivers and streams further mirrors this calculation^39^. The lower solubility of high elevations indicate both higher ebullition rates and contribution to the overall methane flux according to current ebullition models which utilize an equilibrium concentration threshold of methane to predict^44^. In lowland areas, sporadic high ebullition events can be triggered by weather events or changes in pressure gradients when a critical gas volume threshold is exceeded^32,33^. By assuming a constant mass of methane for a given bubble, the gas expands and methane concentration becomes diluted along rising elevation (as described by Equation 4, Methods), and this lead to a reduced methane concentration in bubbles but a more rapid gas ebullitions (Table S2). This is evidenced by our comparative analysis of methane concentrations in bubbles from high-elevation sites versus those from lower elevations (Table 1). While these data support our interpretation that bubbles become more dilute but more voluminous at high elevations, the limited sample size (particularly for mid-elevations) constrains the quantification of bubble dynamics across the full gradient. Expanded bubble sampling across diverse mountain regions would strengthen the linkage between theoretical modelling and field observation.

Our findings indicate that the decrease in pressure with increasing elevation exerts two primary effects that enhance methane ebullition. Firstly, there is a ‘degas’ or ‘outgas’ effect, which is a consequence of decreased solubility and significantly promotes the formation of methane-containing bubbles. This is analogous to the “saturation-controlled regime” observed during water table fluctuations in deep water bodies^19^. Secondly, there is a ‘trigger’ effect that increases the volume of bubbles^17^, potentially enabling their upward propagation to overcome the resistance of the pore structure^45^. The close agreement between the theoretical model (derived solely from the pressure-dependent ‘degas’ and ‘trigger’ effects) and the empirical trend across the full elevation gradient provides strong evidence that these two physical mechanisms are the primary drivers of enhanced methane ebullition in high-elevation shallow waters. Quantitatively, the model explains over 99% of the variance in the empirical trend, indicating that the pressure-driven physical effects capture the dominant signal. This finding bridges the gap between fundamental physical laws and field observations, establishing a mechanistic basis for understanding methane emissions from mountain aquatic ecosystems.

However, we cannot rule out non-linear responses that might become apparent with more complete sampling across the full elevational gradient as currently data is deficient in the mid to high elevations (e.g. 1000 to 3000 m a.s.l.). Similarly, the maxima deviation between the theoretical model and the empirical trend around 2000 m a.s.l. (Table 2) has indicated the unknown mechanism of the pressure-ebullition relationships. Studies from lower elevations have shown that soil porosity can restrict bubble movement, and a critical pressure threshold exists at which the bubble volume surpasses the pore capacity^32,45^. This implies a substantial reservoir of methane-containing bubbles within the soil^46,47^, where the free-phase methane in these bubbles may be re-dissolved or oxidized^44^, or otherwise escape once they overcome the critical pressure threshold.

### Two distinct ebullition modes

Bubble movements in sediments under varying hydrostatic pressures have been delineated into two distinct scenarios: ‘stable’ and ‘dynamic’^48^. In the “stable” scenario, bubble growth under high hydrostatic pressure (>1000 hPa) acts as a counterforce to ebullition, as methane within the bubbles can re-dissolve during their ascent. Conversely, in the ‘dynamic’ scenario, bubble growth under low hydrostatic pressure (< 1 hPa) significantly facilitates the upward movement of bubbles. It is evident that in high-elevation environments, achieving ‘saturation’ for bubble formation is a crucial and advantageous condition for initiating the ‘dynamic’ movement of bubbles.

We recognize the significant spatial variability in sediment porosity, even at the site-specific level^45,46^. However, for the purpose of conceptual clarity, we assume uniform porosity across the elevations under study. Under this assumption, we can delineate two primary modes of methane ebullition that differentiate low from high elevation settings (Fig. 4). At high elevations, there is a greater presence of free-phase methane in the pore water at any given time, a result of ‘degas’ effect, and the total volume of methane bubbles is further increased by ‘trigger’ effect. These two effects are synergistic and collectively augment the facilitation of ebullition-related processes along elevation gradients, though the specifics of key processes, such as the formation and movement of individual bubbles, remain to be elucidated.

**Figure 4.**
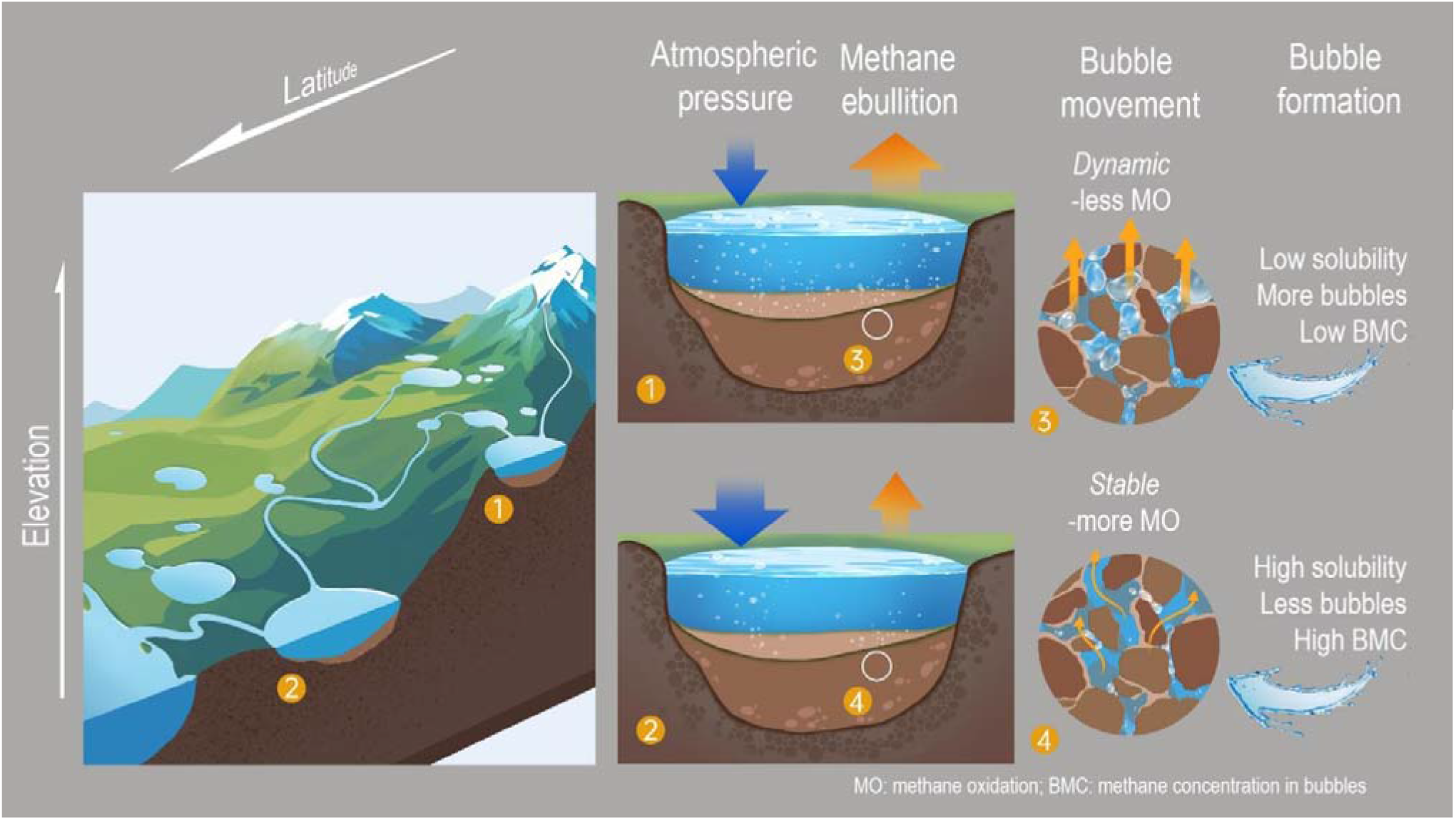
The conceptual model showing different mechanisms controls the methane ebullition from shallow waters at low and high elevations.

Field data comparisons on ebullition characteristics have provided the evidence for distinct ebullition modes at varying elevations. Both the occurrence and the relative contribution of ebullition were observed to escalate with elevation (Table 1). In addition to the underwater video documentation provided in this study (Movie S1), persistent ebullition phenomena have been reported in high-elevation environments by other studies^24,25^. The trend of increasing ebullition with elevation can also be encapsulated alternatively as follows: with rising elevation, a growing proportion of methane produced evades oxidation and is emitted via ebullition, as methanotrophy has been demonstrated to be commensurate with methanogenesis^49,50^. This can be reversely mirrored by the mitigated overall emissions observed in condition that the facilitative effects of low pressure on ebullition were not present^51^.

### A hidden accelerant of global warming

Our findings demonstrate that methane ebullition escalates with elevation due to pressure-dependent ‘degas’ and ’trigger’ effects. This elevation gradient creates a positive feedback loop: disproportion rising trend of temperatures along elevation^52^ amplify methane production in sediments, while declining atmospheric pressure further accelerates methane ebullition. Moreover, permafrost thaw and glacial melt in high mountains could destabilize sediment, facilitating ebullitions, and triggering abrupt emissions as what has been found in Arctic region^24,53^. Furthermore, the data presented in this study predominantly capture the mean ebullition rates during the ice-free season, thereby precluding an analysis of the variations linked to local weather events. As per Equation 4 (see Methods), ebullition is anticipated to exhibit heightened sensitivity to temporal atmospheric pressure changes with increasing elevation (Fig. 3b). Currently, high-resolution spatiotemporal data remain critically scarce, hampering the accuracy of global greenhouse gas budgets^54^. The elevation-dependent patterns uncovered here not only pinpoint priority regions and ecosystem types for future monitoring but also add a previously unrecognized physical dimension to our understanding of methane dynamics.

Our findings pose significant challenges to the current efforts on evaluating the global methane budget (including the tropical mountains that are not mentioned in the present study), as there are not only lakes but also marshes, swamps, streams *etc.* widely distributed across global mountains^55^. In fact, the current global methane emission inventories are still far from complete, with the most important source of uncertainty being natural emissions, especially those from wetlands and other inland waters^1^. This incomplete information distorts current climate policy toward fossil fuel, and waste management, *etc.* for methane mitigation. For instance, Methane Alert and Response System of International Methane Emissions Observation (IMEO), covers only 10 to 17 % of the already known global emissions. When the elevation-driven ebullition is considered, urgent action could be taken to avert a self-reinforcing climate feedback, particularly in vulnerable regions like the Himalayas and Andes. We thus call for: () Elevation-specific monitoring by deploying low-cost bubble traps and establishing monitoring networks for mountain aquatic ecosystems, particularly these with shallow water area, to track ebullition; () Nationally Determined Contributions (NDCs) integration by encouraging countries with significant mountainous regions (e.g., India, Canada, China, United States, and the Andean countries) to incorporate high-elevation water bodies into their NDCs; ()

Nature-based solutions by stabilizing and reducing ebullition in mountain wetlands through restoration, leveraging their dual role as carbon regulators and biodiversity reservoirs. We urge the Intergovernmental Panel on Climate Change (IPCC) to classify mountain aquatic ecosystems as “emerging methane sources” to align science with global mitigation policy.

## Acknowledgments

We dedicate this work to the memory of Binod Shrestha, who contributed to field sampling in Nepal before his untimely passing. We thank the Ministry of Forest and Environment, Government of Nepal, for permitting field research in the national park. We acknowledge Y. J. Dai for fieldwork support with bubble traps, and T. L. Ma and J. Hu for laboratory sample analysis. We are grateful to F. Li for his expert guidance on modeling bubble volume dynamics under reduced atmospheric pressure, and to Y. C. Xia and Y. Gao for figure preparation.

## Author contributions

D.Z., H.B.J., H.C., and N.W. designed the study; D.Z., I.R., K.W.A. and M. K. collected literature data and provided ecological context and interpretation; D.Z., H.B.J., J.L.L., and X.Z. performed, aided in or supervised data analysis and modelling; D.Z., S.J., N.B., and S.J.T. performed field sampling and laboratory experiment; D.Z., H.B.J., and I.R. wrote the manuscript with input from all other authors.

## Data, code, and materials availability

All data are available in the main text or the supplementary materials.

## Methods

### Identification of climate zone

Methane emissions, including ebullition, vary among climate zones. The rate of methane emissions from tropical ecosystems was significantly higher than that from the rest of the globe. Ecosystem productivity and climate are the key regulators for this variation^4,6^. On a global scale, most mountain ecosystems fall into the range of subtropical, temperate, boreal, and sub-arctic zones. The climatic zone of our field sites belong to temperate (Dhaap Lake) and cold temperate (Lake and ponds on the eastern Tibetan Plateau). Generally, the spatial distribution of the study sites is within the permafrost zone of the northern hemisphere (Figure 1a).

### Source of dataset

Ebullition data, along with other related parameters from reference studies^22,24,29–31^ are listed in Table S1. We used water depth (<4 m), sampling period (four months in growing season, except for West Siberia lakes with two months sampling) as the filter to extract related ebullition data from these reference studies. Atmospheric pressures (AT) of each site were calculated from corresponding elevation (EL) (by equation: AT= EL*-9.0811+98905), and hydrostatic pressure (HP) of each site were calculated from water depth (WD) (by equation: HP =WD*101325/10.34).

For those ebullition data associated with the temporal variation^32,33^, a baseline atmospheric pressure has been identified (based on its altitude). The corresponding atmospheric pressure to target ebullition was calculated by deducting the pressure drop (the pressure head were converted to equivalent atmospheric pressure) (Extended Data Fig. 3).

### Field sampling and parameters of ebullition

The Dhaap Lake (2089 m a.s.l.), located at the mid-hill of Nepal, was selected as one of the field study sites (Fig. S1 a). Methane fluxes were measured using the vented closed chamber and gas chromatograph method^60^. Gas sampling was carried out between 09:00 and 12:00 hours in Nepal standard time (GMT+5:45). For each plot, four gas samples from the chamber air headspace were taken into 5 ml airtight vacuumed vials at 0, 10, 20 and 30 min after deployment. Methane emissions were calculated based on the temporal variation of methane concentration in the headspace during the sampling period. The emissions were considered as diffusive when the *R*^2^ of linear correlation between methane concentration and the elapsed time was greater than 0.90, while the emissions were considered as the mix of ebullition and diffusive when *R*^2^ was under 0.90, which indicates one or more abrupt increases of the headspace methane concentration was observed^32^. Due to the difficulty to identify the contribution of diffusive to overall flux for individual chamber when ebullition happens, we defined the average flux rate from chambers without ebullition as the baseline diffusive flux in each sampling campaign. Therefore, the ebullition rate of individual chamber was calculated by subtracting the baseline diffusive flux from the overall flux. We used data from eight sampling campaigns during growing season. The seasonal variation of methane emission from Dhaap Lake is shown in Extended Data Fig. 2.

Three ponds (On the watershed of the Yangtze river and the Yellow river on the southern edge of Zoige Plateau) and one small, shallow lake (Fairy Lake on the northern edge of Zoige Plateau) in the same region of Lake Medo, which was studied in our previous research^22^, was selected as the site for the ebullition survey using bubble traps (Fig. S1 bc). We deployed traps (0.4 m in diameter) that freely floated in the open water area to evaluate the “background” ebullition^41^. On August 7 and 8, 2022, we deployed six and five traps in the ponds (two for each) and the lake, respectively. The trapped air samples were measured for their volumes (extremely fast bubble accumulation was observed, i.e., 8 to 189 ml of bubble samples were trapped within approximate two to five hours, Table S2) and collected with 10 ml vacuum vials in the field. The samples were then taken back to the laboratory for methane concentration analysis. The methane concentrations in the bubbles are shown in Table 1 and Table S2.

### Modelling methane solubility and pressure

The relationship of methane solubility (*c,mol/m*^3^) versus water temperature and pressure could be described as follows^35^:

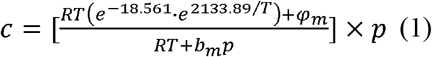

in which

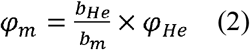

and in which

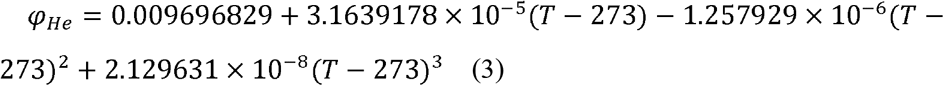

Where *c* is the methane solubility in *mol/m*^3^,*_m_* is the Van der Waals volume of the methane, which is 4.28 × 10^−5^*m*^3^/*mol*, the value of 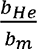 is calculated as 0.4058. is the absolute pressure in Pa, *T* is the gas temperature in *K*, and *R* is the universal gas constant. The methane solubility controlled by atmospheric pressure (altitude) and temperature could therefore be quantified, and a three dimension figure (Fig. 3a) showing their relationship has been generated by Matlab (R2020a) accordingly (see Table S2).

### Modeling bubble volume enlargement

According to mass balance, a single gas bubble in liquid phase follows:

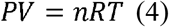

Where *V* is the volume of gas bubble, *n* is moles of gas bubble, and *T* is the absolute temperature in the bubble, respectively, and *R* is the universal gas constant. *P* is the total pressure and its value should be a sum of atmospheric pressure (*P*) and hydrostatic pressure (*pwgh*, where *pw* is the mass density of the fluid phase, *g* is the gravitational acceleration, *h* is depth of the bubble center), therefore the volume of bubble at an initial phase *(V_0_*) can be given as:

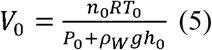

When a change in atmospheric pressure (Δ) occurs along temporal scale, regardless of its temporal or spatial variation, a condition shift, named phase *t*, allows us to compute the volume of bubble (*Vt*) as:

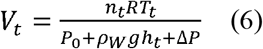

Therefore, the variation of gas bubble volume is:

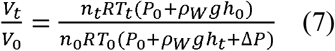

We assume that the mass, temperature, and depth of the gas bubble do not change along with atmospheric pressure variation; they are regarded as constant in our modeling. Thus, equation (4) can be simplified as:

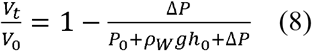

If P_0_ = α P_s_; ρwgh_0_ = β *P*_s_ Δ P= γ *P*_s_ when *P*_s_ as standard atmospheric pressure was introduced, then the equation can be given as:

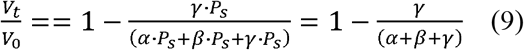

In conditions where αɛ[0.46546,1] (equivalent to altitude from 0 to 6000 m a.s.l.), γ ɛ[0.00, 0.04] (equivalent to an atmospheric pressure variation of 40 hPa), and a depth of 0.5 m water column for bubble, i.e.β =1, the volume change of gas bubble controlled by atmospheric pressure in both spatio-temporal scales could be quantified, and a three dimension figure (Fig. 3b) showing their relationship has been generated by Matlab (R2020a) accordingly (see Table S3).

### Comparison between the theoretical and empirical ebullition

We developed a physical model that integrates the ‘degas’ and ‘trigger’ effects. According to Henry’s law, the equilibrium solubility (*c*) of methane in pore water decreases linearly with decreasing total pressure (*P*) (at constant temperature). The ‘degas’ effect thus promotes bubble nucleation, and the efficiency of bubble formation (η*degas*) can be expressed as proportional to the inverse of solubility, i.e. η*degas*□1/*c*□1/*P*. For the ‘trigger’ effect, the volume (*V*) of a bubble containing a fixed mass of gas follows the ideal gas law, *V*□1/*P*. Since the ability of a bubble to overcome sediment pore resistance increases with its volume, the release probability (η*trigger*) can be approximated as η*trigger*□*V□*1/*P*. Assuming that methane production (per unit area per unit time) is constant along the elevational gradient (a premise supported by the absence of a corresponding trend in diffusive fluxes (Extended Data Fig. 4)), and as the ‘degas’ effect and the ‘trigger’ effect are independent and sequential, the theoretical ebullition flux is proportional to the product of their efficiencies, yielding *F*□*P*^-2^.

To assess the consistency between the model and the empirical trend, we compared the relative flux generated by the linear empirical model fitted to the field data (Fig. 2b; *F*=-0.3754*P*+484.63) with the theoretical relative flux (derived from the combined ‘degas’ and ‘trigger’ effects) (Fig. 3ab, Table S2, S3 at the temperature of 12°C and temporal variation of pressure at zero) at 500 m elevation intervals. The two fluxes were both normalized to the sea-level value (0 m) with a 0.5 m water column.

## Statistical analysis

All statistical analyses were performed in R software (v.4.4.2). A one-way analysis of variance and Tukey’s least significant difference (LSD) test were employed to assess the significant differences in methane emission rate from the shallow waters (both ebullition and diffusive) between different pressure gradients. Linear regression models were used to examine the relationships between methane ebullition and pressure. A Pearson correlation analysis was used to test the relationship between theoretical relative flux and the empirical relative flux. Visualization of results was carried out using the “ggplot2” and “ggpubr” packages.

**Extended Data Figure 1 to 4**

**Extended Data Table 1**

**Extended Data Figure 1.**
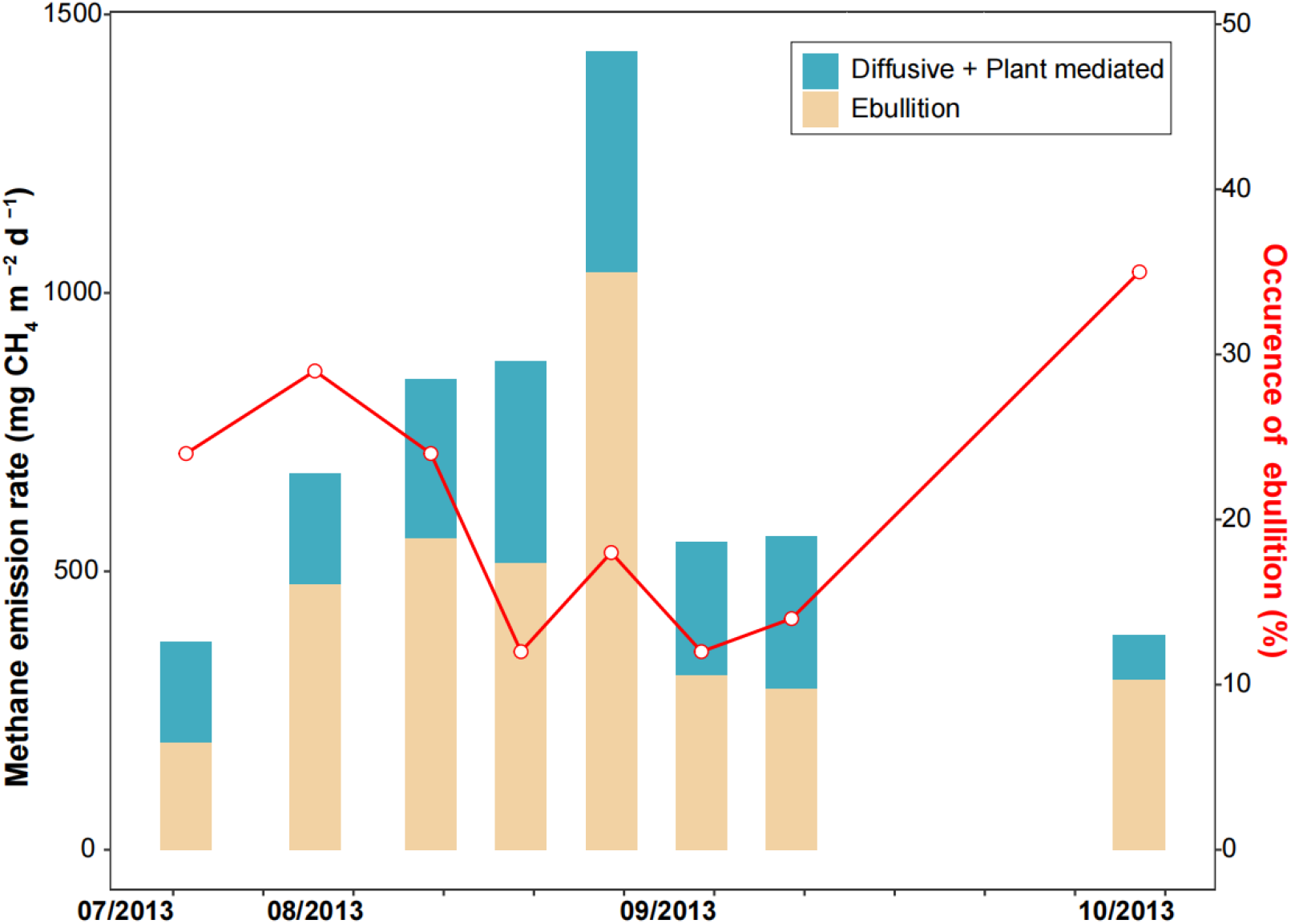
Temporal variation of methane ebullition rate, flux rate of diffusive plus plant mediated and occurrence of ebullition from Dhaap Lake during July to October, 2013.

**Extended Data Figure 2.**
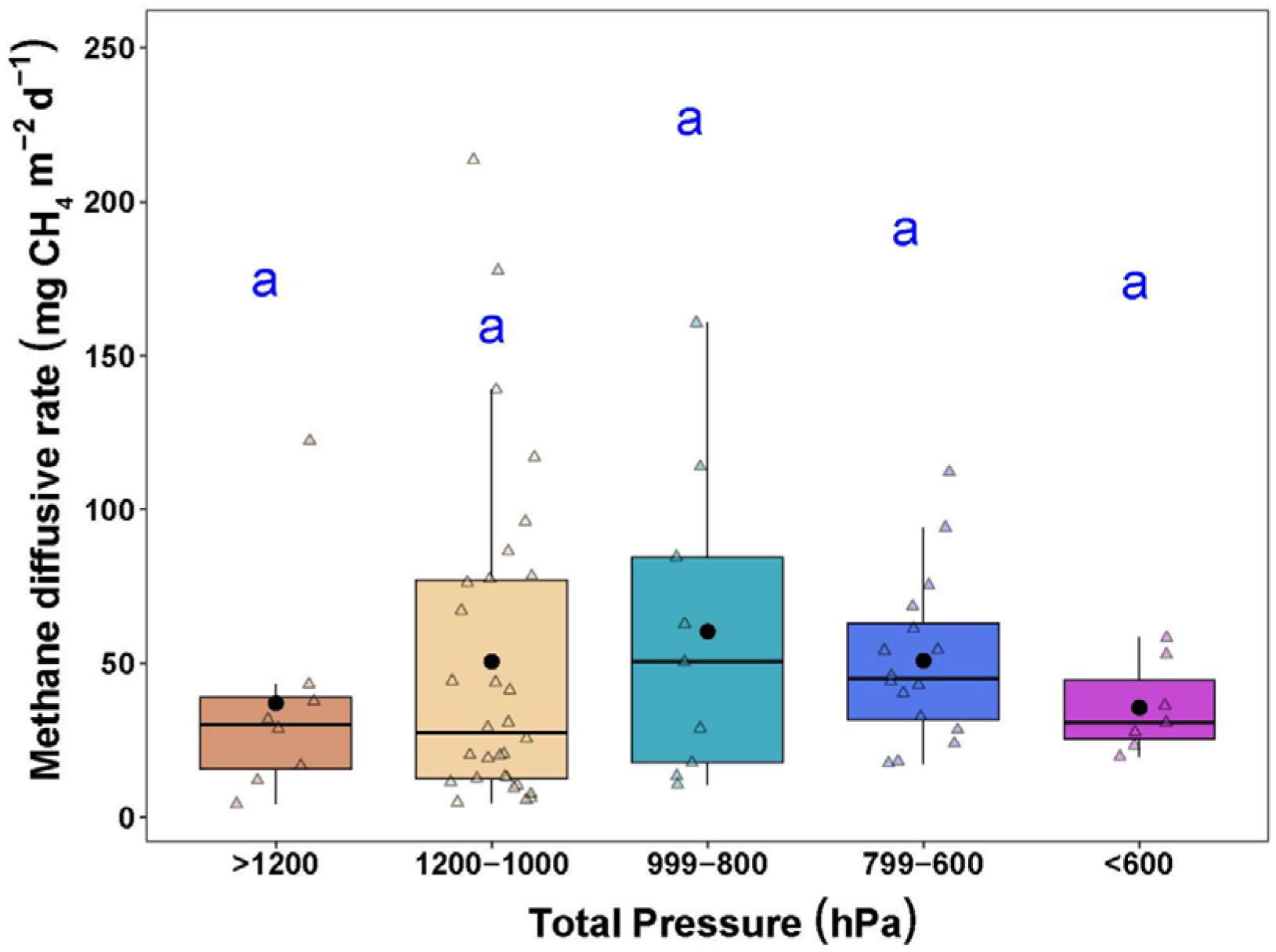
Comparison of the diffusive flux rates by grouped total pressures. Data presented are first quartiles, medians and third quartiles with standard errors.

**Extended Data Figure 3.**
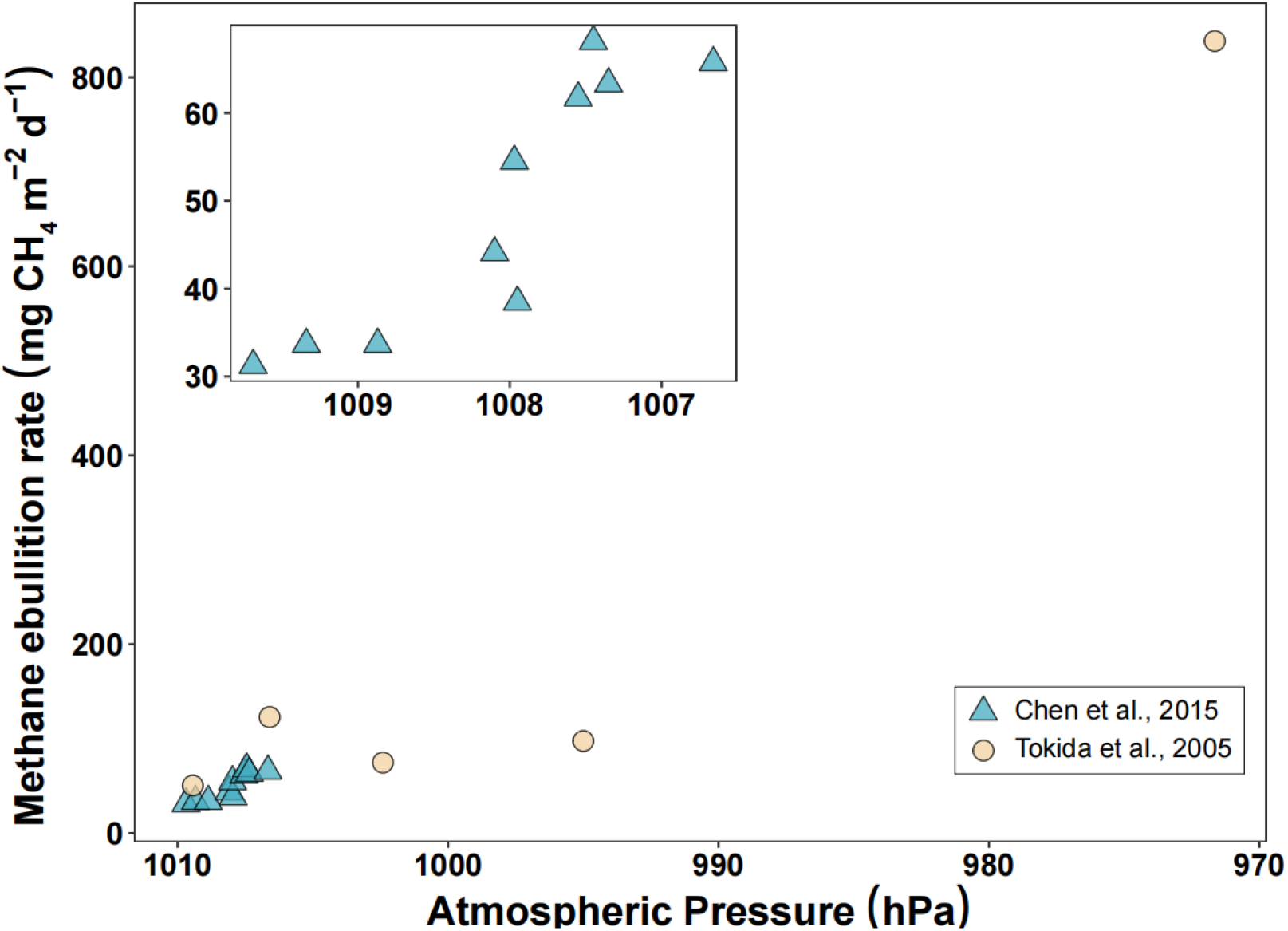
Scatters of methane ebullition rates and atmospheric pressure along time (data source are shown in the lower-right box).

**Extended Data Figure 4.**
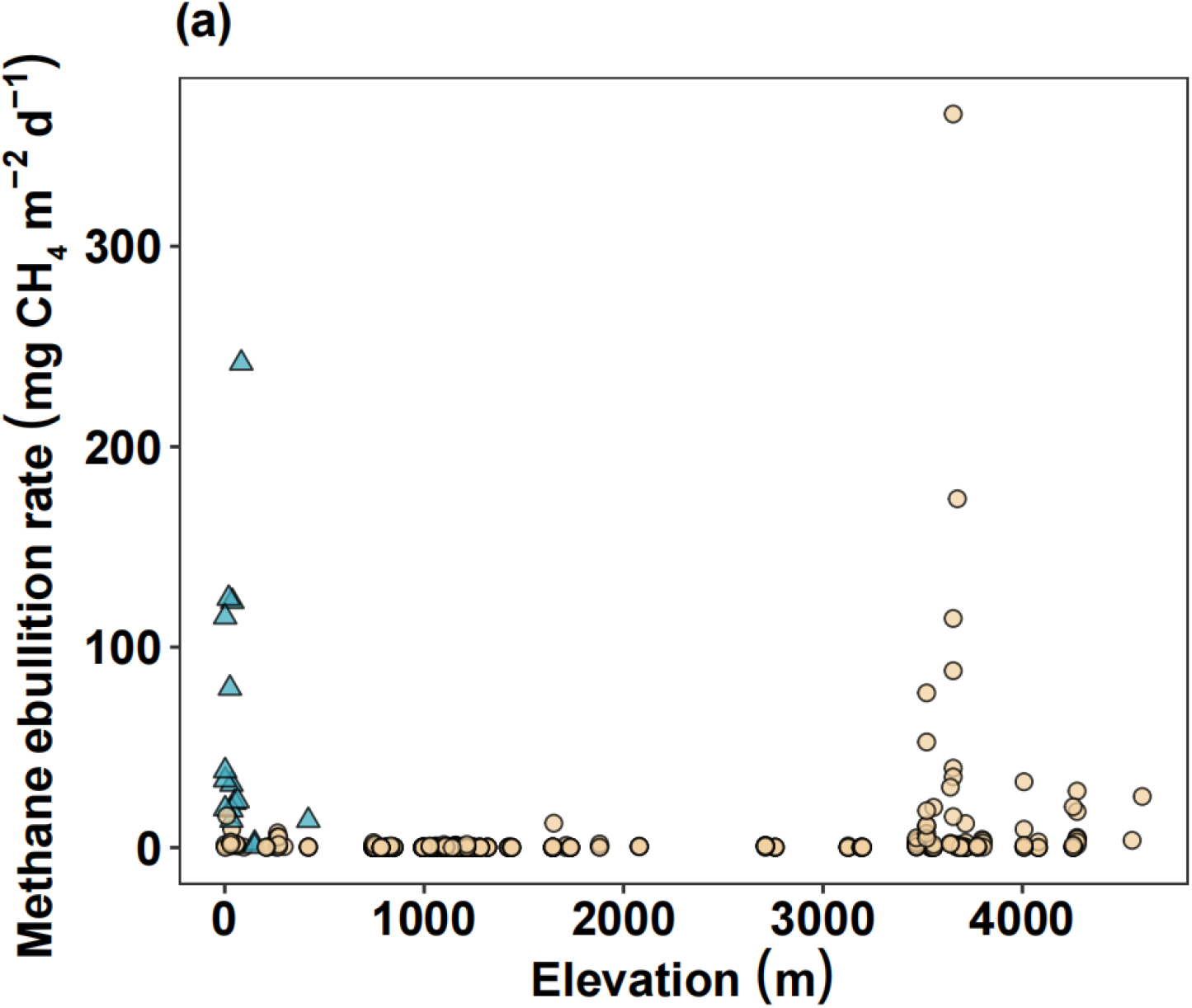
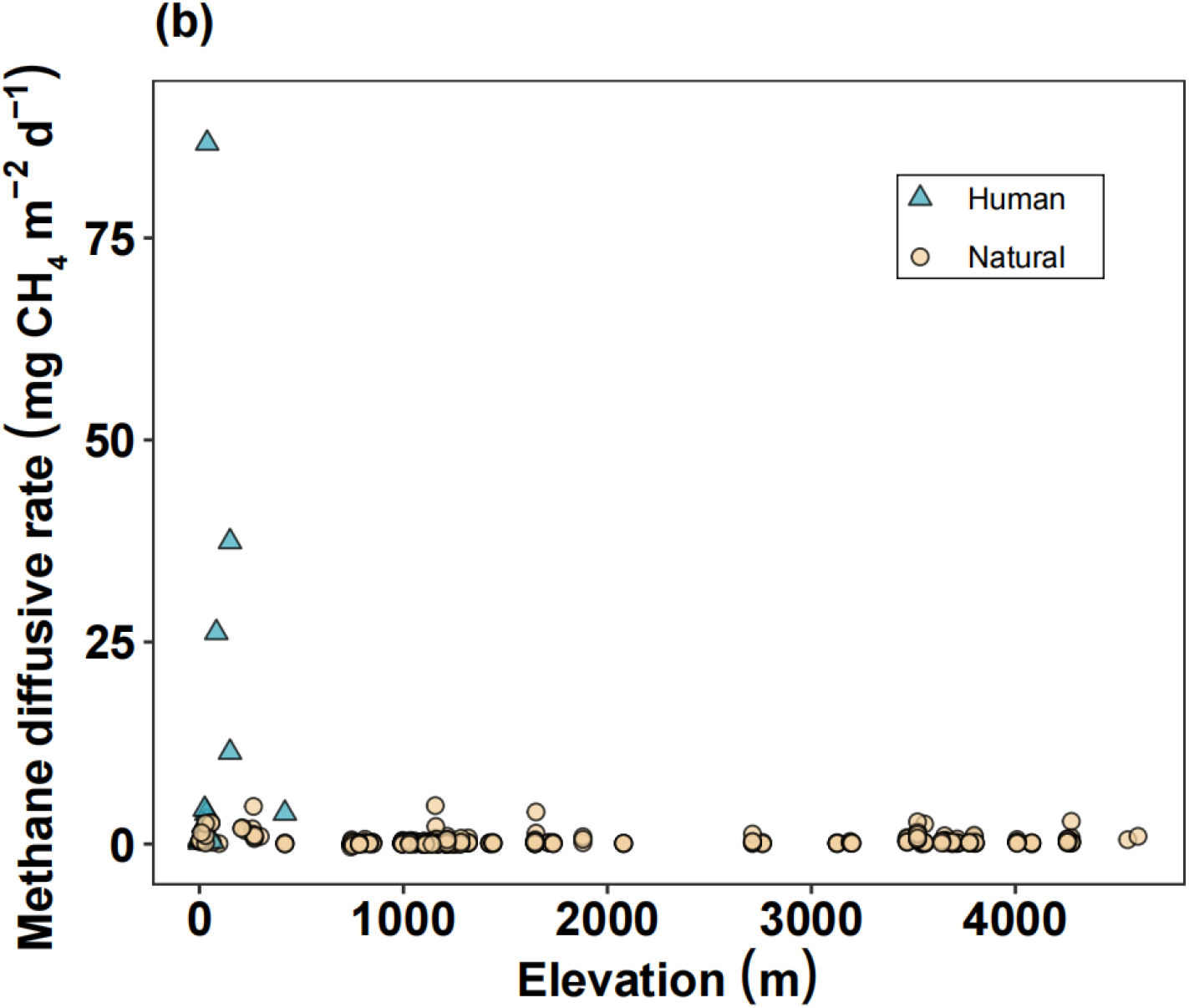
Ebullition (a) and diffusive (b) fluxes of methane against elevation from the the Global River Methane Database (GRiMeDB). High methane ebullition and diffusive rates were found in low elevation rivers which were under intensified human activities (blue triangles in both a and b), while high ebullition rates were observed in high elevations where high diffusive flux were absent for these under natural conditions (yellow round in both a and b). Data in the figure was extracted according to the same criteria in Methods (exclude the sites in tropical zone, sampling period represents for the growing season) and available in Table S4.

**Extended Data Table 1.**
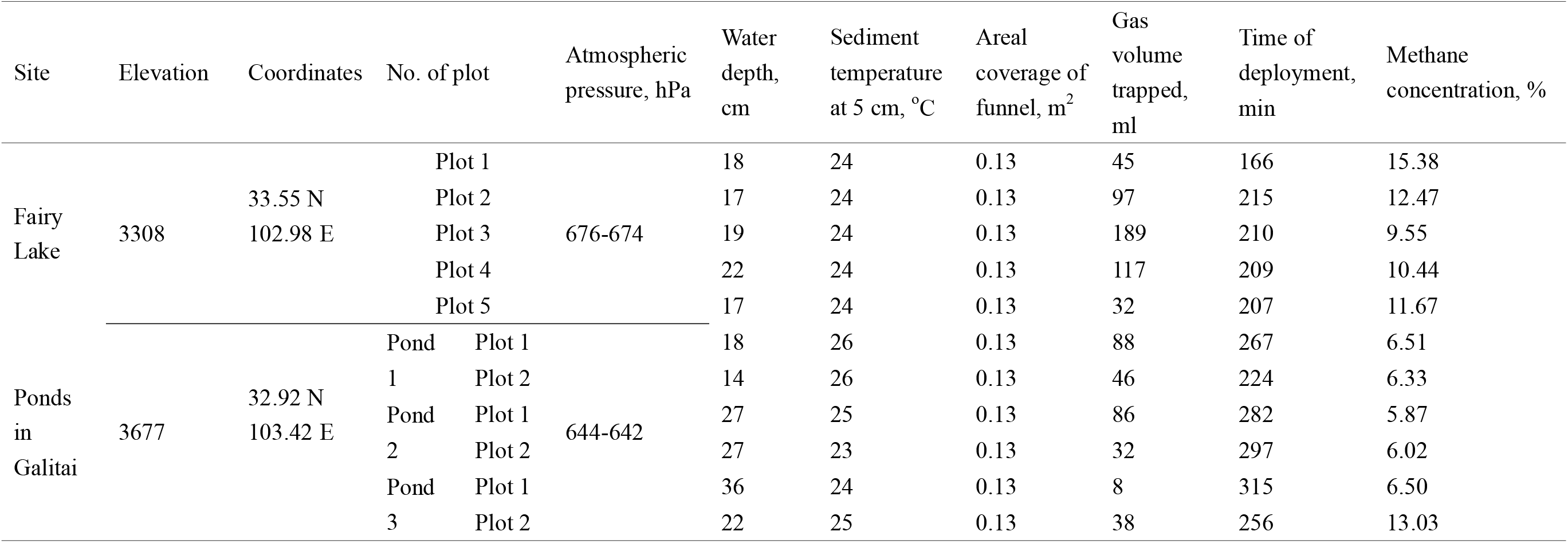
Environmental factors and parameters of bubble traps on the Zoige Plateau.

