## Supplemental Fig. 1 for "Physical laws predict methane hotspots in global mountain waters"

**This file includes:**

Figure S1

Table S1 to S4 which are all in Excel format.

**Other Supplementary Information for this manuscript include the following:**

Movie S1 Persistent ebullition recorded by an underwater camera in a high elevation lake (Fairy Lake).

**Figure S1 The field survey and sampling sites for methane ebullitions in this study. (a)** The Dhaap lake, which locates at an elevation of 2089 m a.s.l., act as one of the headwaters for the Kathmandu Valley of Nepal. **(b)** The ponds scattered on the watershed (Galitai) of the Yangtze river and the Yellow river on the southern edge of Zoige Plateau. **(c)** The Fairy Lake on the northern edge of Zoige Plateau, with five bubble traps deployed during the sampling campaign.

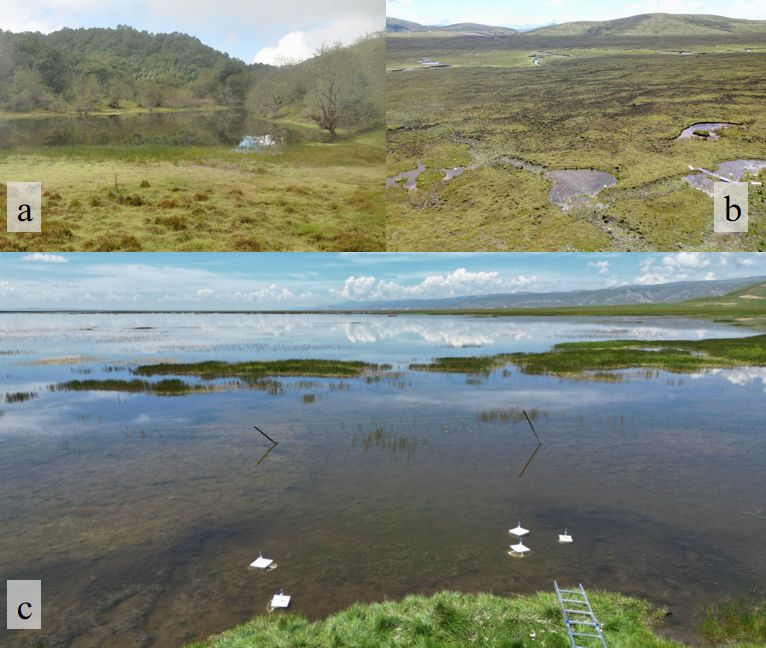

**Table S1 The original data of all the shallow waters in this study.**

Note: Lake type: PP-peatland pond, GP-glacial/post-glacial pond, T-thermokarst pond; Lake bottom: P-peat, O-organic, U-unspecified, Y-yedoma, M-minerogenic; Ebullition measurement method: BT-Bubble trap, CH-Chamber, IS-Ice survey; Diffusive measurement method: CH-Chamber, WS-Water Sample.

a: the diffusive flux cannot be calculated directly because the chamber method involves plant mediated flux as well, thus we assume that the diffusive accounts for 25% of the sum of diffusive and plant mediated flux according to Ref. 67; nd: no data.

**Table S2 Data of the modeled relationship between methane solubility versus atmospheric pressure and temperature.**

**Table S3 Data of the modeled relationship between gas volume and atmospheric pressure varies in both temporal and elevational scale.**

**Table S4 Ebullition and diffusive fluxes data of methane and related parameters from the the Global River Methane Database (GRiMeDB).**
